# Spatial and temporal habitat availability declines towards and beyond the geographic range limit of a coastal dune endemic

**DOI:** 10.64898/2026.03.30.715381

**Authors:** Graydon J. Gillies, Michael P. Dungey, Christopher G. Eckert

**Author notes:** Correspondence author: Graydon J. Gillies Current address: Department of Geography, Memorial University of Newfoundland, 232 Elizabeth Ave, St. John’s, NL A1B 3X9.

## Abstract

1. Changes in habitat structure across species’ distributions may contribute to the generation and maintenance of range limits, but few studies have evaluated this by directly measuring habitat availability across relevant spatial scales.
2. Here, we test the predictions that coarse-scale and patch-level habitat availability decline towards and beyond the northern range limit of Pacific coastal dune endemic *Camissoniopsis cheiranthifolia*. We used aerial imagery and geographic information system (GIS) tools to measure the coarse-scale availability of coastal dune habitat in California and Oregon. The availability of finer-scale habitat patches specifically suitable for *C. cheiranthifolia* was measured in a 2-generation field survey of > 4,200 5m x 5m plots randomly distributed across 1100 km of coastal dune habitat transcending the species northern range limit. At each plot, we estimated the proportion of area that contained suitable habitat as well as recorded occupancy by *C. cheiranthifolia*. As an alternative approach to visually estimating habitat suitability, we recorded plant community composition at each plot to predict beyond-range habitat suitability using a random forest model.
3. Contrary to our predictions, we found that coastal dune habitat, measured coarsely from aerial imagery, was more abundant and continuous towards and beyond the northern range limit. At the fine scale, however, the proportion of plots with suitable habitat (patch suitability) and the proportion of habitat within plots that was suitable (patch size) declined across the range limit. Moreover, patches were more isolated from one another and, in one survey year, less temporally stable towards and beyond the range limit. Finally, occupancy by *C. cheiranthifolia* was less likely in smaller, more isolated, and temporally unstable patches, providing mechanistic insight to the previously observed decline in occupancy towards the range limit.
4. **Synthesis**: Taken together, our results suggest that fine-scale habitat patch configuration changes in ways that likely impede patch colonization, thereby reducing occupancy and limiting the species’ northern distribution. Thus, consideration of geographic variation in patch and landscape structure, rather than only coarse-scale habitat availability, may be essential for understanding the processes that limit species ranges.

## INTRODUCTION

Diverse evolutionary and ecological explanations for species’ geographic distribution limits have been proposed, but few have been adequately tested (Sexton et al., 2009). Yet, predicting how species will respond to global change and, in particular, whether they are able to shift their ranges to track climate (Chen et al., 2011; Tomiolo & Ward, 2018) requires understanding the processes that maintain species range limits. The ecological causes of species range limits tend to be classified into two general categories. A species may be unable to expand its range because the habitat beyond is not suitable for population self-replacement (“niche limitation”).

Alternatively, suitable habitat may occur beyond the range limit but the species lacks the capacity to successfully disperse to it (“dispersal limitation”). The niche-limitation hypothesis has been tested using beyond-range transplant experiments, but results are mixed. Individual fitness often declines when species are moved beyond their range limits, but not enough to prevent populations from persisting (Brown et al., 1996; Hargreaves et al., 2014; Lee-Yaw et al., 2016). When the niche limitation hypothesis fails, dispersal limitation is often invoked as an untested default (though there is some supporting evidence in a few case studies; see Gaylord & Gaines, 2000). “Absolute” dispersal limitation occurs where there is some major obstacle to dispersal or a gap in suitable habitat preventing species from accessing suitable beyond-range habitat. An alternative and more nuanced form of dispersal limitation, inspired by metapopulation theory, is that range limits may be imposed by subtle declines in habitat connectivity in a manner that impedes the patch-to-patch colonization required to balance local extinction of the species from patches. This can, in theory, generate an abrupt range limit where the occupancy of suitable habitat falls to zero (the “metapopulation hypothesis” for range limits; Carter & Prince, 1981; Holt & Keitt, 2000).

This metapopulation hypothesis for range limits suggests that a range limit (i.e., zero patch occupancy) can be imposed when patch colonization declines and/or patch extinction increases towards the edge of a species’ geographic distribution (Carter & Prince, 1981; Holt & Keitt, 2000; Lennon et al., 1997), but this has almost never been tested. In theory, range limits can be caused by relatively subtle changes in one or both parameters, and both parameters can be affected by habitat characteristics. For instance, colonization should decline as patches of suitable habitat become less frequent on the landscape, smaller in size, or more isolated from each other by unsuitable habitat (Pellet et al., 2007). Conversely, extinction partly reflects limits on survival and reproduction imposed by a species’ niche (Grinder & Wiens, 2023). The likelihood of patch extinction will increase as habitat quality and, consequently, individual fitness declines. However, there is also a stochastic component to patch extinction not caused by variation in habitat quality but instead by disturbance or succession (i.e., patch turnover; Griffen & Drake, 2008; Reigada et al., 2015). As a consequence of unfavourable shifts in metapopulation dynamics, occupancy of suitable habitat patches should decline. In a review of empirical studies on geographic variation in demography and population structure, Pironon et al. (2017) found that occupancy of habitat declined towards species’ range margins in 81% of tests whereas declines in fitness were much less consistent (declining population growth in 32% of tests; declining fecundity in 43%), suggesting that geographical variation in metapopulation dynamics, especially colonization, might commonly contribute to the maintenance of range limits.

However, it remains unclear why exactly habitat occupancy declines. It is likely that geographic variation in habitat structural characteristics directly mediate processes of habitat colonization and extinction, but few studies have comprehensively quantified geographic variation in habitat structure towards range limits, especially over ecologically relevant spatial scales. Thus, quantifying habitat patch characteristics such as frequency, size, isolation, and turnover across large scales is important for evaluating the underlying mechanistic causes that dictate geographic variation in colonization and extinction rates towards species range limits.

Although variation in habitat patch structure across species range limits has almost never been extensively quantified, there is a substantial body of work on metapopulation biology outside the context of species distributions, particularly for plants. However, there is little consistency in which parameters these studies find are important for dictating metapopulation dynamics. For example, incorporating environmental features (e.g., vegetation cover, grazing rates, and adjacent habitat types) in a metapopulation model did not predict patch occupancy of the butterfly *Melitaea cinxia* better than a model that included only patch size and isolation (Moilanen & Hanski, 1998). In contrast, habitat quality but not patch size or isolation predicted patch occupancy by the perennial plant *Colchicum autumnale* (Adriaens et al., 2009). While it is evident that the structure of fragmented habitat can influence habitat occupancy, no study to our knowledge has comprehensively quantified multiple components of habitat structure towards species range limits. Furthermore, studies that examine variation in habitat structure towards and beyond range limits tend to examine habitat availability at coarser spatial scales, and not in conjunction with fine-scale measures of habitat structure such as patch isolation or temporal stability. Thomas et al. (1999) used data on habitat availability for four heathland animals (silver-studded blue butterfly, a red ant, heathland grasshopper, and sand lizard), measured from vegetation composition, land-use, and topographical variables, and found that available habitat patches became less frequent towards their range limits, but they did not quantify other measures of habitat structure. Similarly, Angert et al. (2018) used ecological niche modelling to show that estimated habitat suitability and occupancy declined towards latitudinal and elevational range edges of *Erythranthe cardinalis*, but this coarse-scale analysis did not directly examine geographic variation in other fine-scale patch characteristics such as size or isolation.

While patch characteristics can evidently vary spatially, they also vary temporally (Reigada et al., 2015). Occupancy of suitable patches can be transient as subpopulations go extinct due to demographic stochasticity, but the suitability of patches themselves can also shift between states of suitable and unsuitable. Temporal variation in habitat suitability may be predictable and repeatable, such as temporary ponds that provide spring habitat for aquatic species before drying out (Ferreira et al., 2015; Jeffries, 2008), or it may be more stochastic, for instance when unpredictable and infrequent desert rains provide ephemerally suitable habitat for annual plants (Venable, 2007). Habitat may also be made temporarily unsuitable through natural disturbances (Wilcox et al., 2006). Or, suitable habitat for more opportunistic ruderal species may transition to unsuitable habitat through succession. While the turnover of suitable habitat by definition increases extinction rates in metapopulations (Vuilleumier et al., 2007; Wilcox et al., 2006), opportunistic, early-successional species might have higher occupancy under moderate patch disturbance as they quickly colonize disturbed habitat patches before late-successional species make patches unsuitable again (Mestre et al., 2020). However, to our knowledge, there are no empirical studies that attempt to quantify how temporal variation in habitat availability changes towards species range limits (but see Benning et al., 2022 for a modelling approach). Should disturbance of suitable patches become more frequent towards a range limit, occupancy may decline (or perhaps increase for opportunistic species). Thus, understanding temporal dynamics in habitat structure across space is important for understanding how species occupy their environment.

Here, we use the endemic Pacific coastal dune plant *Camissoniopsis cheiranthifolia* (Hornem. Ex Spreng.) W.L. Wagner & Hoch (Onagraceae) to test the hypothesis that habitat structure changes towards and beyond its northern range edge in ways that contribute to reducing patch occupancy and maintaining its range limit. Replicated transplant experiments with *C. cheiranthifolia* have shown that the species can persist with high fitness for multiple generations well beyond its northern range limit, suggesting niche limitation alone is not responsible for maintaining this range limit (Cross & Eckert, 2021; Samis et al., 2016; Samis & Eckert, 2009). Absolute dispersal limitation also seems unlikely because coastal dune habitat extends hundreds of km beyond the northern range limit, although geographic variation in the availability of dune habitat has not been quantified until now. Gillies et al. (2025) previously used a large-scale cross-generation habitat survey to show that the occupancy of suitable habitat patches declines toward the northern range limit and this is entirely accounted for by declining patch colonization rates.

However, it remains unclear how habitat characteristics might vary towards the range limit in a manner that could inhibit dispersal and reduce colonization rates. They showed that colonization was less likely in smaller patches and with lower local abundance (i.e., lower propagule pressure), and both patch size and local abundance declined towards the range limit (Gillies et al., 2025). However, it remains unclear how other habitat characteristics (such as patch isolation and turnover) vary towards the species northern geographic range limit. Furthermore, how each of these habitat characteristics change beyond the northern range limit where the metapopulation is unable to persist despite the presence of suitable habitat (Cross & Eckert, 2021) remains untested. Here, we evaluate how colonization of beyond-range habitat might be impeded by measuring multiple habitat characteristics both towards and beyond the species northern range limit.

Coastal dune habitat tends to be highly fragmented and prone to regular disturbance (Austrich et al., 2021; Malavasi et al., 2018) and thus is ideal for investigating geographic variation in both coarse-scale habitat availability and fine-scale habitat patch dynamics. Here, we assess how habitat structure changes across the range edge using two approaches. First, we measured geographic variation in coastal dune habitat availability, regardless of suitability for *Camissoniopsis cheiranthifolia*, coarsely defined using aerial imagery. Second, we surveyed the finer-scale spatial and temporal variation in the availability of habitat patches specifically suitable for *C. cheiranthifolia* within coastal dune habitat over two generations. We measured habitat suitability, occupancy, and plant community composition in ∼4,100 5m x 5m plots spanning ∼1100 kilometres of coastline towards and beyond the species’ northern range limit, using randomly sampled plots as a proxy for habitat patches. We predict that, towards and beyond the species range limit:

1. The availability and/or connectivity of coastal dune habitat, coarsely defined with aerial imagery, declines, which may impose absolute dispersal limitation on the species’ northern distribution.
2. The availability of habitat patches specifically suitable for *C. cheiranthifolia* within coastal dune habitat, measured as the proportion of plots that contain suitable habitat (patch suitability) from visual surveys, and as predicted by plot-level plant community composition, declines.
3. The area within plots that contains suitable habitat (patch size) declines.
4. The distance between plots with suitable habitat (patch isolation) increases.
5. Patches become less temporally stable (i.e., greater patch turnover/lower patch stability).
6. Occupancy of patches by *C. cheiranthifolia* is more likely in patches that are larger, less isolated from other suitable patches, and more temporally stable.

## MATERIALS AND METHODS

### Study system

*Camissoniopsis cheiranthifolia* is endemic to coastal dune habitat from northern Baja California, Mexico to central Oregon, USA. The species occurs exclusively on relatively stabilized dune below 50 m elevation with coarse silt to medium sand and low organic matter soils that are subject to moderate disturbance by wind and water to keep the density of competing vegetation low (Samis & Eckert, 2007). Thus, suitable habitat for *C. cheiranthifolia* is highly fragmented by areas of unsuitable habitat including severely disturbed dune, highly vegetated dune, standing water, and driftwood accumulation. This species also has a particularly short generation time of 1.02–1.92 years (Samis et al., 2016). Like many coastal dune endemics, *C. cheiranthifolia*’s distribution stretches more than a thousand kilometres of latitude but no more than a few kilometres inland from the coast (McLachlan, 1991), meaning its geographic distribution is near one-dimensional and virtually linear. Its small seeds (0.2 mg) seem wind- and water-dispersed (Dungey, 2021), which suggests dispersal likely occurs along this “one-dimensional” coastal habitat.

### Coastal dune habitat structure

For analyses that test geographic variation in both coarse and fine-scale habitat metrics towards the range limit, we measured the distance to the range limit along the coast as the predictor by generating a “coastal reference line” that traces the coastline using the Iso Cluster Unsupervised Classification tool in ArcMap software (v. 10.8; see Dungey, 2021; Gillies et al., 2025) which better reflects the distance along which *C. cheiranthifolia* disperses than distance calculated linearly “as the crow flies”. To assess geographic variation in the availability of coastal dune habitat, coarsely defined, we first used aerial imagery to generate 92,167 points at 10-m intervals along the 1117 km coastal reference line from San Francisco, California to Pacific City, Oregon. We then classified these points as either dune habitat or non-dune habitat (using criteria in Dungey, 2021). To evaluate the availability of coastal dune habitat, we binned these points into 37 groups of 2500 points and calculated the proportion of points in each bin that occurred in or directly adjacent to dune habitat. We also evaluated the fragmentation of coastal dune habitat by measuring the size of the gaps (in metres) between contiguous stretches of coastal dune habitat as the length of continuous runs of one or more points classified as non-dune habitat.

### Suitable habitat patch structure

To test geographic variation in fine-scale habitat patches specifically suitable for *C. cheiranthifolia* towards the species’ range limit, we first delineated shapefiles of putative coastal dune habitat from Google Earth images of the coastline between San Francisco, CA to Pacific City, OR. We then generated 7,031 random point coordinates at about constant density within these shapefiles to survey on foot in 2019 and 2022 (Dungey, 2021; Gillies et al., 2025). We located (in 2019) and relocated (in 2022) each point with a handheld GPS (Garmin GPSmap 60Cx) and established a 5m x 5m plot centered on each point. We verified that plot centers could be relocated with little error (∼ 1m, see Gillies et al., 2025). Within each plot, we first classified the habitat as either coastal dune habitat or non-dune habitat, where the latter was open and highly disturbed beach habitat, standing water or anthropogenic cover such as pavement. Plots with no coastal dune habitat were not analyzed further, leaving 4386 dune plots that were analyzed in 2019 and 4265 in 2022. For plots containing dune habitat, we first classified whether a plot contained any habitat specifically suitable for *C. cheiranthifolia* (hereafter “patch suitability”) and then estimated proportion of the plot area that contained suitable habitat (then used to calculate “patch size” as the suitable area, m^2^) based on habitat suitability criteria described by Samis & Eckert (2007) and Gillies et al. (2025). We considered habitat unsuitable for *C. cheiranthifolia* if it contained highly wind-blown disturbed sand, was thickly vegetated, or was covered by driftwood, rock, or other debris. Our measures of habitat suitability specific to *C. cheiranthifolia* come from multiple large-scale studies across the species’ geographic distribution (Gillies et al., 2025; Samis & Eckert, 2007, 2009) as well as within- and beyond-range transplant experiments where transplanted individuals enjoyed high fitness in habitat identified as suitable based on these measures (Cross & Eckert, 2021, 2024). While our measure of patch size may underestimate the size of patches greater than 25 m^2^, this approach provides an adequate index of the amount of habitat availability directly at the plot locations. Furthermore, only 1.07% (in 2019) and 5.86% (in 2022) of patches reached the maximum patch size of 25 m^2^ (Fig. S1a). We calculated “patch isolation” for suitable plots as the average straight-line distance (“as the crow flies”) to the five nearest plots that contained some suitable habitat (Fig. S1b). While we attempted to randomly distribute points into coastal dune habitat at constant density, the randomization algorithm resulted in points being at slightly higher density towards the range limit (Dungey, 2021). Thus, any geographic increase in patch isolation towards the range limit, as predicted, would reflect geographic patterns in habitat structure above and beyond any negative effect caused by the distribution of sampling plots. We calculated “patch stability” as a binary variable to reflect the temporal stability of suitable patches, where 1 indicates “stable” plots that contained suitable habitat in both survey years and 0 indicates “unstable” plots that contained suitable habitat in only one of two survey years. Thus, the group of “stable” plots were suitable in both years (and thus designated as “stable” in both years). Plots deemed unstable for the 2019 analysis were suitable in 2019 and unsuitable in 2022. Plots deemed unstable for the 2022 analysis were unsuitable in 2019 and suitable in 2022. While it is possible that patches transitioned between states of suitable and unsuitable in the time between each survey, this metric provides at least an approximate means of quantifying the temporal stability of each habitat patch. We also recorded occupancy by *C. cheiranthifolia* in each plot.

Changes in plant community composition often reflect changes in relevant environmental conditions (Diekmann, 2003) and thus habitat suitability (Sawchik et al., 2003). To that end, in 2022, we measured the plant community composition at each plot by recording all plant species that occurred on the 5m x 5m cross centred in the plot (Table S2). We identified 107 species within the range limit of *C. cheiranthifolia*, and 85 of these species occurred in more than one plot. As an alternative approach to estimating the quantity of suitable habitat towards and beyond the species’ range limit, we trained a random forest model to predict occupancy both within and beyond the range limit as a proxy for the abundance of suitable habitat (below).

### Analyses

We conducted all analyses in R (v. 4.5.1; R Core Team, 2025) running in RStudio software (v. 2025.09.1+401; Posit team, 2025). We evaluated prediction 1, that the availability of coastal dune habitat, coarsely defined, declines towards and beyond the northern range limit, by fitting variation in the proportion of points in each of the 37 spatial bins that occurred in coastal dune habitat to a generalized linear model (GLM) with coastal distance to the range limit as the single predictor and the beta-binomial error distribution to account for overdispersion. We fit this and all following models using the *glmmTMB* function from the glmmTMB package in R (v. 1.1.11; Brooks et al., 2017). We evaluated the significance of this and all subsequent predictors using type II tests performed by the *Anova* function from the car package (v. 3.1-3; Fox & Weisberg, 2019). For each model that used the distance to the range limit as a predictor, 0 represents the northernmost plot at which we observed *C. cheiranthifolia*, positive values represent plots situated towards the range core (up to +938 km south of the range limit), while negative values represent plots situated north of the range limit (up to –178 km), hence positive regression coefficients indicate an increase in response variables southward to the range core. We again used the *glmmTMB* function to fit variation in inter-dune gap sizes (the lengths of the gaps between coastal dune sites, using each gap as an independent observation) to a linear model with distance to the range limit as the sole predictor.

To evaluate geographic variation in the structure of habitat specific to *C. cheiranthifolia*, we evaluated how patch suitability, size, isolation and turnover varied towards the range edge (predictions 2 to 5). Because each survey involved a slightly different set of plots (see above), the analyses described below were performed on the 2019 and 2022 data separately. First, we tested prediction 2 that patch suitability declines towards the range limit by first fitting binary patch suitability to a binomial GLM with the distance to the range limit as the sole predictor. We then tested prediction 3 that patch size (m^2^) declines towards the range limit and prediction 4 that the log_10_-transformed patch isolation increases towards the range limit by fitting each to a Gaussian GLM with the distance to the range limit as the predictor, again using the *glmmTMB* function. Both the patch size and patch isolation models excluded plots that were entirely unsuitable (patch size of zero) to avoid confounding with variation in patch suitability. We tested prediction 5 that patch stability declines (i.e., patches have lower temporal availability) towards the range limit by fitting binary patch stability to a binomial GLM as above. For each patch stability model, we fit the models using only patches that were suitable in each respective year. Sample sizes for each model are generally lower than the total number of dune plots observed (Table 1) as we were unable to collect all data at some plots (e.g., for some plots we estimated binary suitability but could not estimate patch size) and some models are fit using only a subset of all plots (e.g., our patch size model was fit using only patches > 0 m^2^).

**Table 1.** Binomial and Gaussian GLM analyses of the effect of range position (distance to the northernrange limit) on multiple characteristics of patches suitable for coastal dune endemic *Camissoniopsis cheiranthifolia*. All models are fit using the 2019 or 2022 survey data separately. Positive coefficients represent a decline in the response variable towards the range limit. Patch characteristics that are significantly related to range position are bolded.

| Response | Year | df | *n* | *b* | $\chi^2 / F$ | *P* |
| --- | --- | --- | --- | --- | --- | --- |
| Patch suitability | **2019** | **1** | **4386** | **0.0014 log-odds** | **180.79** | **<0.0001** |
| (Binomial) | **2022** | **1** | **4118** | **0.0010 log-odds** | **87.59** | **<0.0001** |
| Patch size | 2019 | 1 | 1384 | 6.54x10<sup>-4</sup> | 1.37 | 0.29 |
| (Gaussian) | **2022** | **1** | **2089** | **4.24x10<sup>-3</sup>** | **41.99** | **<0.0001** |
| log<sub>10</sub>(Patch isolation) | **2019** | **1** | **1383** | **-1.94x10<sup>-4</sup>** | **39.01** | **<0.0001** |
| (Gaussian) | **2022** | **1** | **2089** | **-1.28x10<sup>-4</sup>** | **25.88** | **<0.0001** |
| Patch stability | 2019 | 1 | 1565 | -6.24x10<sup>-5</sup> log-odds | 0.064 | 0.80 |
| (Binomial) | **2022** | **1** | **2088** | **1.19x10<sup>-3</sup> log-odds** | **50.30** | **<0.0001** |

As an alternative approach to our visual estimates of patch suitability (prediction 2), we used plot-level plant community data in a random forest (RF) model to predict the plot-level occupancy of *C. cheiranthifolia* in plots both within and beyond the range. RF models were chosen because of their ability to incorporate a large number of predictors and their interactions to yield output classifications with high accuracy (Cutler et al., 2007). We fit a RF model using all within-range coastal dune plots, regardless of suitability status and occupancy by *C. cheiranthifolia*, where the binary occupancy of *C. cheiranthifolia* is predicted by the binary occupancy of all species observed within the range (i.e., excluding eight species that were only observed north of the range limit; details in Appendix S2). The random forest model had an out-of-bag classification error rate (an estimate of the error rate based on predictions of samples not used in the bootstrap used for a given tree in the RF model; Janitza & Hornung, 2018) of 16.97%, with most of this error attributed to poor identification of occupied patches (256/693 occupied patches classified correctly; 63.06% class error rate) rather than unoccupied patches (2421/2531 unoccupied patches classified correctly; 4.34% class error rate). Weighting the model classes by the inverse proportion of their frequency in the training data decreased the error rate of patch occupancy classifications to 6.78% but substantially increased the unoccupied classification error rate to 52.39% and more than doubled the overall out-of-bag error rate to 42.59%, so we proceeded using the unweighted model. We then used this RF model to make out-of-bag predictions of *C. cheiranthifolia* occupancy at each plot throughout the entire study region, both within and beyond the range. We then fit variation in the model-predicted *C. cheiranthifolia* occupancy to a binomial GLM with the distance to the range limit as the sole predictor. To assess the importance of each species for prediction in the RF model, we reported the percent increase in mean standard error (MSE) when each species is removed from the model.

Lastly, we assessed prediction 6 that occupancy by *C. cheiranthifolia* is more likely in plots that are larger, less isolated, and more stable. We fit variation in occupancy as a binary response variable to a GLM with patch size, log_10_-transformed patch isolation, and binary patch stability as predictors. Patch size and isolation were scaled to a mean of zero and SD of 1, so that partial regression coefficients could be compared among predictors and years.

We checked that the residuals from all linear models conform to model assumptions with diagnostic plots, tested for homoscedasticity and influential outliers, and evaluated whether GLMs fit model assumptions using tools in the DHARMa package (v. 0.4.7; Hartig, 2022).

## RESULTS

### Coastal dune habitat structure

Across 1117 km of coast from San Francisco, CA to Pacific City, OR the proportion of points that occurred in coastal dune habitat varied from 0% (at the Lost Coast, California where coastal dune habitat is exceedingly rare) to 95.6% near Winchester Bay, Oregon (mean ± SD = 42.9 ± 30.3%). Contrary to prediction 1, the proportion of points that occurred on dune sites increased towards the northern range limit (*n* = 37 coastal segments, *b* = –2.07x10^−3^, χ*^2^* = 8.55, df = 1, *P* = 0.0034; Fig. 1a). This pattern holds when all coastal segments without any dune habitat (i.e., the Lost Coast) are excluded (*n* = 33, *b* = –2.05x10^−3^, χ*^2^* = 14.95, df = 1, *P* = 0.00011; Fig. S2a).

**Figure 1.**
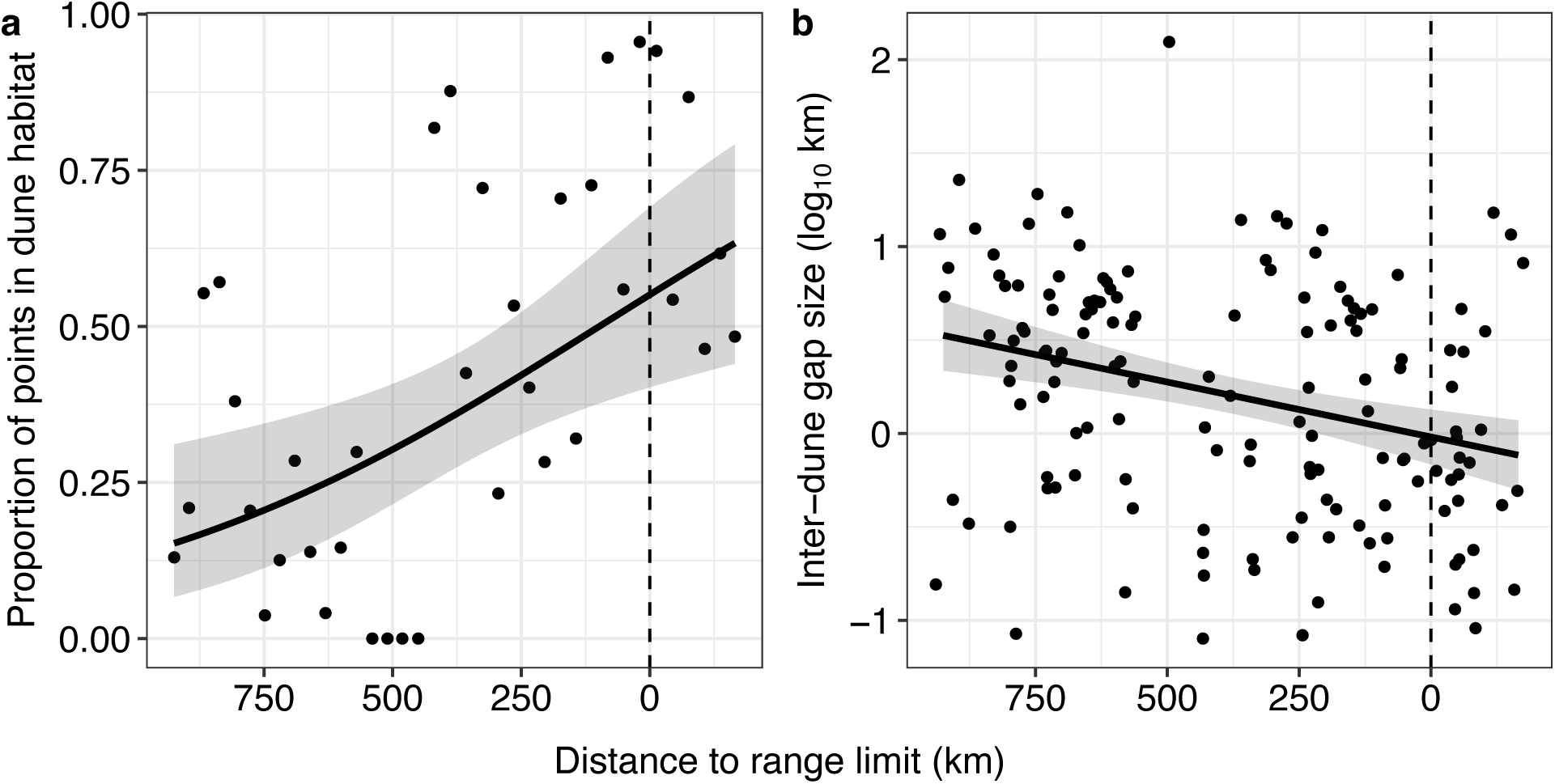
The availability of coastal dune habitat increased and the fragmentation of dune habitat decreased towards and beyond the northern range limit of *Camissoniopsis cheiranthifolia*. The proportion of uniformly spaced points (binned into 37 groups of 2500) increases while the gap distance between adjacent stretches of contiguous coastal dune declines towards and beyond the range limit. Grey ribbons are 95% confidence intervals. The vertical dashed lines indicate the northern range limit (43.8°N).

Coastal dune fragmentation measured as the length of gaps between areas of contiguous coastal dune habitat ranged 80 m to 124.36 km, with a mean ± SD gap size of 4.27 ± 10.75 km. Contrary to prediction 1, the log_10_–transformed gap size declined significantly towards and beyond *C. cheiranthifolia*’s range limit (*n* = 149, *b* = 5.88x10^−4^, χ*^2^* = 15.15, df = 1, *P* < 0.0001; Fig. 1b), even after excluding the large gap that occurs across the Lost Coast (*n* = 148, *b* = 5.74x10^−4^, χ*^2^* = 15.26, df = 1, *P* < 0.0001; Fig. S2b). Notably, the first gap north of the northernmost dune site occupied by *C. cheiranthifolia* was only 550 m, which falls in the lower 23^rd^ percentile of gap sizes that occur within the range, well below the median within-range gap size of 2,429 m.

### Habitat patch structure

Overall, 41.3% and 50.8% of plots contained some habitat deemed specifically suitable for *C. cheiranthifolia* in 2019 and 2022, respectively. Consistent with prediction 2, the frequency of plots that contained suitable habitat (patch suitability) declined significantly towards and beyond the range limit in both years (Fig. 2, Table 1 for all patch structure models). Mean ± 1 SD estimated patch size (excluding plots with no suitable habitat) was 11.76 ± 7.13 m^2^ in 2019 and 12.44 ± 8.60 m^2^ in 2022. Patch size declined significantly towards the range limit in 2022 but not 2019. Isolation of plots with some suitable habitat ranged 37.07 m to 18.20 km (391 ± 1167 m) in 2019 and 21.8 m to 27.30 km (303 ± 982 m) in 2022. Consistent with prediction 4, isolation increased significantly towards the range limit in 2019 and 2022. Somewhat consistent with prediction 5, patch stability declined significantly towards the range limit in 2022 but not in 2019.

**Figure 2.**
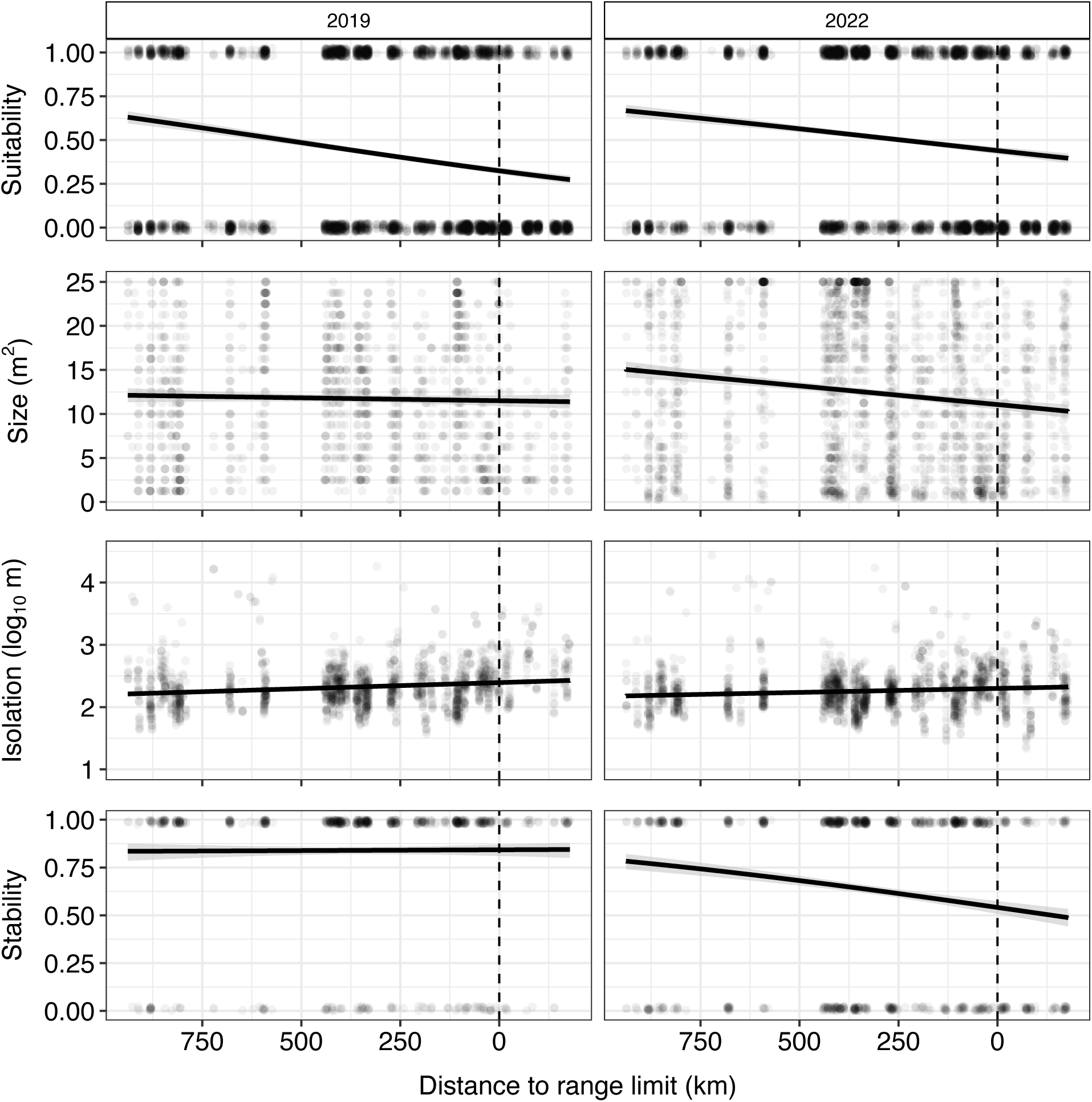
Habitat structure variables (patch suitability, patch size in 2022, patch isolation, and patch stability in 2022) changed significantly towards and beyond the northern range limit of *Camissoniopsis cheiranthifolia*. Horizontal axes depict the distance to the species’ northern range limit. Results from 2019 and 2022 are shown in the upper and lower panels, respectively. Grey ribbons are 95% confidence intervals. Vertical dashed lines indicate the northern range limit (43.8°N). Points in panels (a) and (d) are vertically jittered for clarity.

In our 2022 plot survey, 21.50% of 3223 plots were occupied by *C. cheiranthifolia*. Consistent with prediction 2, the random forests model classified 11.35% (366/3224) of within-range patches as occupied compared to only 0.88% (7/799) of beyond-range patches. Moreover, community-predicted patch occupancy based on random forests analysis declined towards and beyond the species’ northern range limit (*n* = 3919, *b* = 1.87x10^-3^ log odds, χ*^2^*= 113.04, df = 1, *P* < 0.0001; Fig. 3).

**Figure 3.**
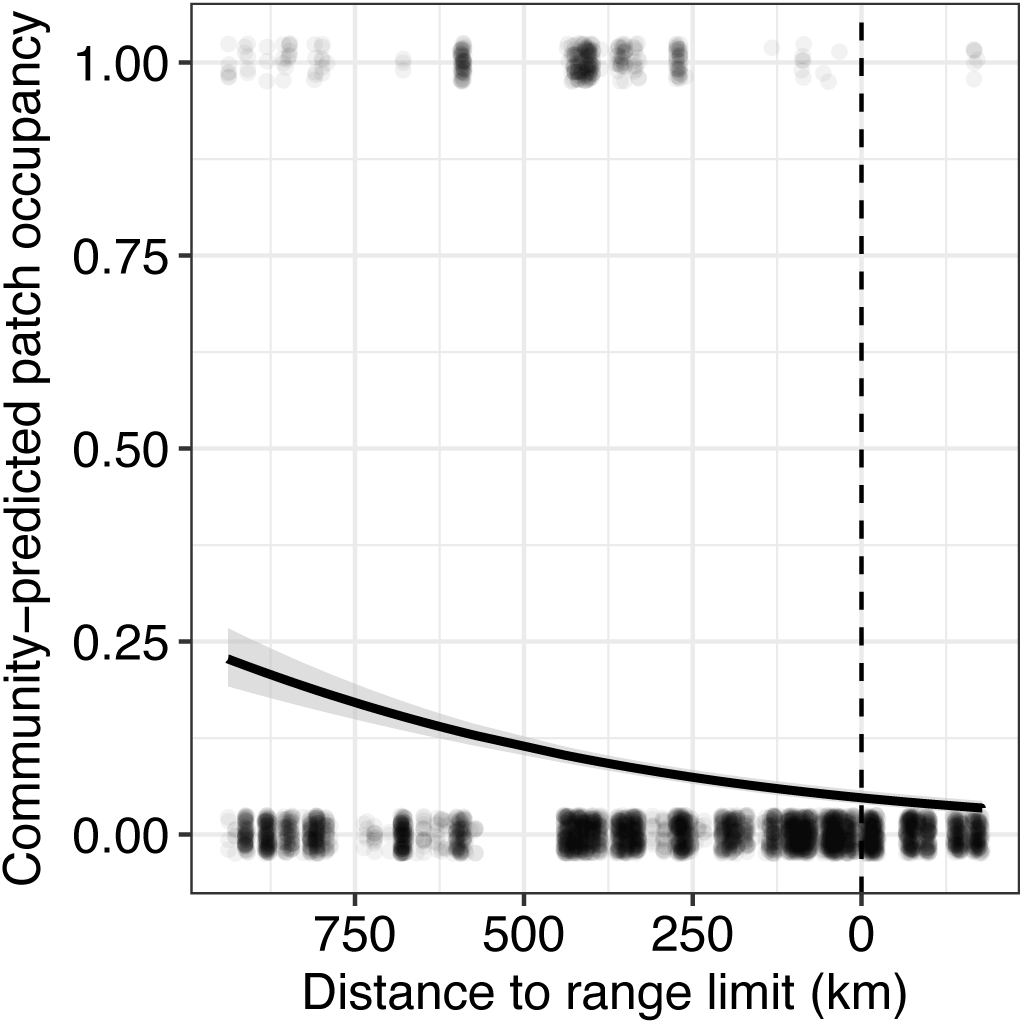
Predicted occupancy of habitat patches declined towards and beyond the northern range limit of *Camissoniopsis cheiranthifoli*a. Occupancy of each plot was predicted by a random forest model trained on occupancy and plant community data within the range. The solid line indicates predictions by a GLM with 95% confidence intervals in grey. The dashed vertical line indicates the northern range limit. Points are jittered vertically for clarity.

### Predicting occupancy

In both 2019 and 2022, patch occupancy increased with patch size and patch stability and decreased with patch isolation, consistent with prediction 6 (Fig. 4, Table 2 for patch occupancy models).

**Figure 4.**
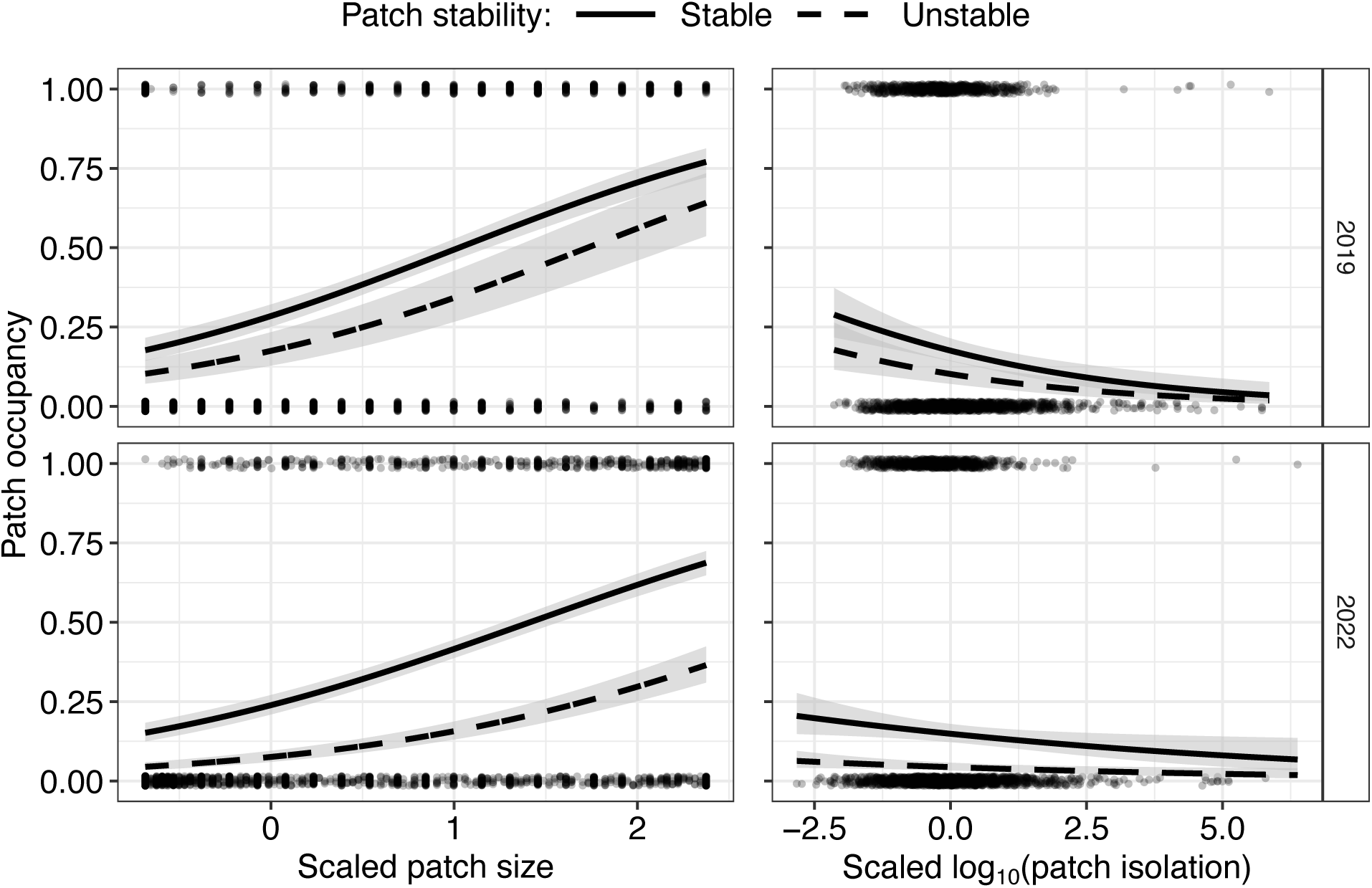
Patch occupancy by *Camissoniopsis cheiranthifolia* increased with patch size (left) and declined with the log_10_-transformed patch isolation (right) in both 2019 (top row) and 2022 (bottom row). Occupancy was also greater in temporally stable patches (solid lines) than temporally unstable patches (dashed lines). Lines show model-predicted occupancy while patch isolation is held constant at its median value (left) and patch size is held constant at its median (right). Grey ribbons are 95% confidence intervals. Points are jittered vertically from 0 and 1.

**Table 2.** Binomial GLM analyses of the effects of patch characteristics on patch occupancy by coastal dune endemic *Camissoniopsis cheiranthifolia* in 2019 and 2022. Patch isolation was log_10_-transformed, then all predictors were scaled for comparability, and all coefficients are in log-odds units. Significant predictors are bolded.

| Response | *n* | Predictor | df | *b* | $\chi^2$ | *P* |
| --- | --- | --- | --- | --- | --- | --- |
| 2019<br>occupancy | 1293 | **Patch size** | **1** | **0.90** | **156.20** | **<0.0001** |
|  |  | **Patch isolation** | **1** | **−0.30** | **19.30** | **<0.0001** |
|  |  | **Patch stability** | **1** | **0.63** | **10.58** | **0.0011** |
| 2022<br>occupancy | 2088 | **Patch size** | **1** | **0.82** | **228.44** | **<0.0001** |
|  |  | **Patch isolation** | **1** | **−0.14** | **5.47** | **0.019** |
|  |  | **Patch stability** | **1** | **1.34** | **116.62** | **<0.0001** |

## DISCUSSION

### Coastal dune habitat structure

Replicated transplant experiments suggest that the northern range limit of *C. cheiranthifolia* is not solely limited by low fitness (Cross & Eckert, 2024; Samis & Eckert, 2009), as experimentally transplanted populations have persisted beyond the range limit for > 10 generations with individual fitness as high as experimental populations within the range (Cross & Eckert 2021). Our test of prediction 1 also suggests that dispersal limitation due to limited or fragmented habitat beyond the edge (“absolute” or “long distance” dispersal limitation) is also unlikely to limit the northern distribution. Coastal dune habitat, coarsely defined from aerial imagery, was more frequent and more continuous beyond the range limit than within the northern portion of range (Fig. 1). The gaps between contiguous stretches of dune at the range limit itself were not large, with the gap distance to the dune site just beyond the northernmost occupied site being only 550 m. The species’ range encompasses much larger gaps that consist of rocky shores and very little coastal dune habitat (e.g. the Lost Coast = ∼124 km, and Big Sur farther south than our survey region, which extends ∼145 km; Walton, 2008) yet the species distribution transcends these major gaps within the range, suggesting that *C. cheiranthifolia* is capable of dispersing much farther than the modest gap in coastal dune habitat found at its northern range limit. Cross & Eckert (2021) found descendant *C. cheiranthifolia* up to ∼500 m from transplanted populations after 10 generations, which suggests that seed can disperse substantial distances over moderate time scales.

Physical barriers to dispersal that involve major gaps in suitable habitat such as mountain ranges, oceans, or rivers are generally thought to be one of the simplest mechanisms that impose range limits (Gaston, 2003). However, it has been argued that, beyond the coarsest of scales (such as dispersal across continents for marine species or across oceans for terrestrial species), such barriers only rarely impose species’ range limits (Merriam, 1894). While there is a growing number of studies that use beyond-range transplant experiments to test for fitness limitation with dispersal limitation as a default alternative (Hargreaves et al., 2014; Lee-Yaw et al., 2016), this approach provides little insight to the nature of the barriers to dispersal or the scale at which dispersal limitation might act. Nevertheless, there is limited evidence for the importance of dispersal limitation shaping species’ distributions. For example, Gaylord & Gaines (2000) showed that ocean currents can inhibit dispersal of pelagic larvae to consequently limit the geographic ranges of marine species. Similarly, the species ranges of Amazonian primates often coincide with major rivers, and several studies found that the similarity of primate species on opposing river banks was lower for wider rivers with greater discharge, suggesting rivers play a role in limiting dispersal (Ayres & Clutton-Brock, 1992; Helenbrook & Valdez, 2025). However, tests of dispersal limitation remain rare, and attributing a species’ range limit specifically to physical barriers to dispersal is challenging. Here, we demonstrated that absolute dispersal limitation caused by suddenly reduced availability and/or increased fragmentation of coastal dune habitat beyond the range likely does not contribute to maintaining the northern range limit of *C. cheiranthifolia*. Instead, dispersal dynamics across habitat patches within coastal habitat are likely far more important.

### Habitat patch structure within coastal dunes

Our analysis of variation in the structure of habitat specific to *C. cheiranthifolia* was motivated by theory suggesting that the dynamics of extinction and colonization among habitat patches on landscapes at finer spatial scales can result in a range limit (Carter & Prince, 1981; Holt & Keitt, 2000; Lennon et al., 1997). In contrast to the near-continuous distribution of coastal dune habitat (coarsely defined), the availability of habitat patches specifically suitable for *C. cheiranthifolia* within the coastal dunes of Oregon and California declined towards and beyond the species’ northern range limit. Our results suggest that patch suitability, size, and stability declined, and patch isolation increased towards and beyond the range limit. This pattern of variation is consistent with metapopulation models that can account for species’ range limits. While other work has used species distribution modelling to estimate the distribution of suitable habitat across species range limits (e.g., Lee-Yaw et al., 2016), particularly in the context of climate-driven range shifts, these models may be limited in their capacity to estimate fine-scale variation in habitat structure that is important for determining local population and metapopulation processes, as these models are usually based on coarse-scale environmental variables rather than fine-scale ecological data that we used here to estimate habitat patch suitability (Littlefield et al., 2019). While there are empirical studies that indicate that metapopulation patch structure can dictate patch occupancy (e.g., Pellet et al., 2007; Schooley & Branch, 2009), to our knowledge there are no studies that have simultaneously measured variation in patch suitability, size, isolation, and turnover towards species range limits, as well as investigated how these habitat patch characteristics might limit patch occupancy towards range limits.

Outside of the range limit context, our finding that occupancy by *C. cheiranthifolia* increased with patch size is supported by other work involving a variety of animal taxa (e.g., Fred & Brommer, 2003; Sahlin & Schroeder, 2010; Shake et al., 2012), however studies on plants are few. Generally, population in larger patches tend to be less prone to extinction and/or larger patches may act as larger targets for colonization, consequently increasing occupancy (Gillies et al., 2025; Pellet et al., 2007). We also found that occupancy by *C. cheiranthifolia* declined with greater patch isolation, as predicted by theory (Hanski, 1994) as well as empirical studies on animal species such as the butterfly *Lycaena helle* across moist meadow habitat (Bauerfeind et al., 2009) and the spittlebug *Neophilaenus albipennis* across its host plant *Brachypodium pinnatum* (Biedermann, 2004). Theory confirms that dispersal can be important for maintaining occupancy of habitat by species and ensuring metapopulation persistence (Johst et al., 2002).

This is supported empirically. For example, marine animals with lower dispersal tend to exhibit greater spatial heterogeneity in abundance around the Channel Islands in California (Reed et al., 2000). Evidently, the spatial structure of habitat (such as increasing patch isolation) can increase the dispersal distance required for colonization (or reduce the strength of rescue effects; Brown & Kodric-Brown, 1977; Eriksson et al., 2014; Hanski, 1998) and consequently reduce the rate of patch occupancy. Similar to our study, Pinto & MacDougall (2010) used a field experiment to demonstrate that dispersal limitation appears to constrain occupancy of habitat-edge patches by the understory plant *Viola praemorsa*. Our study adds to this empirical evidence and goes one step further by showing that spatial variation in multiple metrics of patch structure vary geographically towards a species’ range limit in a manner that could impede colonization of unoccupied patches, and it is possible that these changes in habitat structure become too extreme for *C. cheiranthifolia* to persist on the landscape (Holt & Keitt, 2000).

Furthermore, our patch occupancy models showed that temporally stable plots (i.e., plots with less disturbance or successional turnover) have higher rates of occupancy (Fig. 4), as expected by prediction 6. Intuitively, increasing rates of patch destruction inevitably lead to increased patch extinction rates, directly reducing patch occupancy (Johst et al., 2002; Reigada et al., 2015; Wilcox et al., 2006). However, our finding that *C. cheiranthifolia* tends to occupy temporally stable patches more frequently than temporally unstable patches does not necessarily suggest that the plant experiences greater rates of extinction in temporally unstable patches as they transition from suitable to unsuitable via either disturbance or succession, as the probability of patch extinction does not appear to increase towards this species’ range limit (Gillies et al., 2025). This suggests that, instead, temporally stable patches that are more common at the range core might promote higher occupancy because they are more readily or more frequently colonized by *C. cheiranthifolia*. That is, temporally stable patches may have been available to be colonized for a longer period and thus are more likely to be occupied.

### Causes of declining patch occupancy

Our analyses suggest that geographical variation in patch size, isolation, and temporal stability play at least some role in reducing patch occupancy towards the northern range limit of *C. cheiranthifolia*. However, it is possible that other unaccounted factors might also contribute to declining occupancy. As a post-hoc analysis, we again fit patch occupancy in 2019 and 2022 to binomial GLMs with patch size, isolation, and temporal stability as predictors, but also included distance to the range limit as a covariate, checking the parameter variance inflation factors with the *check_collinearity* function in the performance package (v. 0.16.0; Lüdecke et al., 2021).

Surprisingly, we detected a significant effect of distance to the range limit on patch occupancy above and beyond the effects of patch size, isolation, and temporal stability (Table S1), suggesting other unmeasured factors might also be inhibiting patch occupancy towards the range limit. For example, while our results describe the spatial structure of habitat patches, we are unable to comment on whether or to what extent geographic variation in the quality of habitat patches or the hostility of the habitat matrix between patches might play a role in reducing patch occupancy towards the range limit. To do so would require a carefully designed transplant experiment across habitat patches, thorough demographic surveys, or perhaps measurement of the seedbank within the matrix to quantify the abundance of propagules dispersed into unsuitable habitat.

Alternatively, it is possible that unmeasured intraspecific trait variation towards the range limit might affect successful patch colonization or persistence and consequently patch occupancy. While fitness does not decline towards or beyond the species’ range limit (Cross & Eckert, 2021; Samis & Eckert, 2009) we do not know whether dispersal traits, such as seed size or provisioning, might vary geographically. Theory suggests that increasingly isolated habitat should select for lower dispersal ability due to the increased risk of propagules landing in the inhospitable matrix (Hargreaves & Eckert, 2014; Travis & Dytham, 1999). On the other hand, dispersal ability could also increase where temporal variability in patch availability increases (Hargreaves & Eckert, 2014; McPeek & Holt, 1992), but we only found evidence of declining patch stability in our second survey year. Nonetheless, our results show that while the spatial distribution of suitable habitat patches plays a role in reducing occupancy towards the species’ range limit, there are likely other components of the environment or evolutionary feedbacks that further reduce patch occupancy beyond the direct effects of how habitat is spatially arranged.

### Implications

The occupancy of suitable habitat patches by *C. cheiranthifolia* declines towards the northern range limit, a pattern that can be fully explained by declining inter-patch colonization (Gillies et al., 2025). Here, we have shown that physical habitat structure changes in ways that inhibit occupancy towards and beyond this species’ range limit. Many human activities dramatically alter the key aspects of habitat structure that, based on our results, seem important for maintaining habitat occupancy. For instance, human activities such as urbanization, agriculture and forestry tend to reduce the size and increase the isolation of remnant habitat patches for many native species (Fischer & Lindenmayer, 2007), which may ultimately lead to extinction from landscapes (Grilli et al., 2015; Hanski & Ovaskainen, 2000; White & Smith, 2018). This effect might be most dramatic at the edges of geographic distributions. In many high-latitude countries, the biotically richest areas are in the most equatorial latitudes. For example, most beetles that are red-listed (i.e., threatened) in Finland actually have extensive distributions in other more southerly jurisdictions (Komonen, 2007). In Canada, most at-risk plant species reach their northern range limits in southern Canada but are more widely distributed to the USA farther south (Caissy et al., 2020). If, for some species, range limits are maintained by changes in habitat structure and consequently altered metapopulation dynamics, then conservation efforts to maintain range-edge populations should best focus on the amount and connectivity of suitable habitat regardless of whether that habitat is currently occupied by the species in question.

Unfortunately, these range edges are often where human population density and anthropogenic activity is greatest, creating a perceived conflict between effective conservation and human development (Coristine & Kerr, 2011). Some of these edge populations at risk could be examples of the “living dead”, whereby habitat fragmentation or other human activity has reduced colonization or increased extinction beyond which the metapopulation will remain viable over the long-term (Hanski et al., 1996). The conservation value of geographically peripheral populations has been widely debated but they might be particularly valuable if they are genetically distinct from populations at the range core (Lesica & Allendorf, 1995; Rehm et al., 2015; Van Natto & Eckert, 2022). Conserving and enhancing habitat connectivity at species range edges may be even more important as changing climate forces species range shifts.

## ACKNOWLEDGEMENTS

We thank Elyse Muir, Krista Williamson and Regan Cross for assistance with field surveys; Fran Bonier, Ryan Danby, Steve Lougheed, and Paul Martin for feedback on earlier versions of the manuscript; Karen Samis and Ben Misiuk for helpful discussion; Oregon State Parks, California State Parks, the United States National Forest Service, the US National Parks Service (Point Reyes National Seashore), and the Humboldt Bay National Wildlife Refuge for access to field sites; the Natural Sciences and Engineering Research Council of Canada (NSERC) and Government of Ontario for graduate scholarships to G.J.G.; and NSERC for a Discovery Grant to C.G.E.

## AUTHOR CONTRIBUTIONS

All authors conceived the ideas. MPD conducted the GIS work. All authors designed the field survey and collected data. GJG and MPD conducted the analyses with input from CGE. GJG led the writing of the manuscript with substantial input from CGE; all contributed critically to the drafts and gave final approval for submission.

## DATA AVAILABILITY STATEMENT

Data and code available from Zenodo digital repository: https://doi.org/10.5281/zenodo.19337895

## CONFLICT OF INTEREST STATEMENT

The authors declare no conflict of interest.

## Notes

https://doi.org/10.5281/zenodo.19337895

### Competing Interest Statement

The authors have declared no competing interest.

## REFERENCES

Adriaens, D., Jacquemyn, H., Honnay, O., & Hermy, M. (2009). Conservation of remnant populations of *Colchicum autumnale* – The relative importance of local habitat quality and habitat fragmentation. Acta Oecologica, 35(1), 69–82.

Angert, A. L., Bayly, M., Sheth, S. N., & Paul, J. R. (2018). Testing range-limit hypotheses using range-wide habitat suitability and occupancy for the scarlet monkeyflower (*Erythranthe cardinalis*). The American Naturalist, 191(3), E76–E89.

Austrich, A., Mapelli, F. J., Mora, M. S., & Kittlein, M. J. (2021). Landscape change and associated increase in habitat fragmentation during the last 30 years in coastal sand dunes of Buenos Aires Province, Argentina. Estuaries and Coasts, 44(3), 643–656.

Ayres, J. M., & Clutton-Brock, T. H. (1992). River boundaries and species range size in Amazonian primates. The American Naturalist, 140(3), 531–537.

Bauerfeind, S. S., Theisen, A., & Fischer, K. (2009). Patch occupancy in the endangered butterfly *Lycaena helle* in a fragmented landscape: Effects of habitat quality, patch size and isolation. Journal of Insect Conservation, 13(3), 271–277.

Benning, J. W., Hufbauer, R. A., & Weiss-Lehman, C. (2022). Increasing temporal variance leads to stable species range limits. *Proceedings*. Biological Sciences, 289(1974), 20220202.

Biedermann, R. (2004). Patch occupancy of two hemipterans sharing a common host plant. Journal of Biogeography, 31(7), 1179–1184.

Brooks, M. E., Kristensen, K., van Benthem K, J., Magnusson, A., Berg, C. W., Nielsen, A., Skaug, H. J., Mächler, M., & Bolker, B. M. (2017). GlmmTMB balances speed and flexibility among packages for zero-inflated generalized linear mixed modeling. The R Journal, 9(2), 378.

Brown, J. H., & Kodric-Brown, A. (1977). Turnover rates in insular biogeography: Effect of immigration on extinction. Ecology, 58(2), 445–449.

Brown, J. H., Stevens, G. C., & Kaufman, D. M. (1996). The geographic range: Size, shape, boundaries, and internal structure. Annual Review of Ecology and Systematics, 27, 597–623.

Caissy, P., Klemet-N’Guessan, S., Jackiw, R., Eckert, C. G., & Hargreaves, A. L. (2020). High conservation priority of range-edge plant populations not matched by habitat protection or research effort. Biological Conservation, 249(108732), 108732.

Carter, R. N., & Prince, S. D. (1981). Epidemic models used to explain biogeographical distribution limits. Nature, 293(5834), 644–645.

Chen, I.-C., Hill, J. K., Ohlemüller, R., Roy, D. B., & Thomas, C. D. (2011). Rapid range shifts of species associated with high levels of climate warming. Science, 333(6045), 1024–1026.

Coristine, L. E., & Kerr, J. T. (2011). Habitat loss, climate change, and emerging conservation challenges in Canada. Canadian Journal of Zoology, 89(5), 435–451.

Cross, R. L., & Eckert, C. G. (2021). Long-term persistence of experimental populations beyond a species’ natural range. Ecology, 102(8), e03432.

Cross, R. L., & Eckert, C. G. (2024). Is adaptation associated with long-term persistence beyond a geographic range limit? Evolution, qpae092.

Cutler, D. R., Edwards, T. C., Jr, Beard, K. H., Cutler, A., Hess, K. T., Gibson, J., & Lawler, J. J. (2007). Random forests for classification in ecology. Ecology, 88(11), 2783–2792.

Diekmann, M. (2003). Species indicator values as an important tool in applied plant ecology – a review. Basic and Applied Ecology, 4(6), 493–506.

Dungey, M. P. (2021). A broad scale investigation of dispersal constraints on the northern range limit of a Pacific coastal dune plant (Publication No. 28445132) [Master’s Thesis, Queen’s University]. ProQuest Dissertations & Theses Global.

Eriksson, A., Elías-Wolff, F., Mehlig, B., & Manica, A. (2014). The emergence of the rescue effect from explicit within- and between-patch dynamics in a metapopulation. Proceedings of the Royal Society B: Biological Sciences, 281(1780), 20133127.

Ferreira, F. S., Tabosa, A. B., Gomes, R. B., Martins, F. R., & Matias, L. Q. (2015). Spatiotemporal ecological drivers of an aquatic plant community in a temporary tropical pool. Journal of Arid Environments, 115, 66–72.

Fischer, J., & Lindenmayer, D. B. (2007). Landscape modification and habitat fragmentation: a synthesis. Global Ecology and Biogeography: A Journal of Macroecology, 16(3), 265–280.

Fox, J., & Weisberg, S. (2019). An R Companion to Applied Regression (Third). Sage. https://socialsciences.mcmaster.ca/jfox/Books/Companion/

Fred, M. S., & Brommer, J. E. (2003). Influence of habitat quality and patch size on occupancy and persistence in two populations of the Apollo butterfly (*Parnassius apollo*). Journal of Insect Conservation, 7(2), 85–98.

Gaston, K. J. (2003). The structure and dynamics of geographic ranges. Oxford University Press.

Gaylord, B., & Gaines, S. D. (2000). Temperature or transport? Range limits in marine species mediated solely by flow. The American Naturalist, 155(6), 769–789.

Gillies, G. J., Dungey, M. P., & Eckert, C. G. (2025). Evidence that metapopulation dynamics maintain a species’ range limit. Ecology Letters, 28(5), e70128.

Griffen, B. D., & Drake, J. M. (2008). Effects of habitat quality and size on extinction in experimental populations. Proceedings of the Royal Society B: Biological Sciences, 275(1648), 2251–2256.

Grilli, J., Barabás, G., & Allesina, S. (2015). Metapopulation persistence in random fragmented landscapes. PLoS Computational Biology, 11(5), e1004251.

Grinder, R. M., & Wiens, J. J. (2023). Niche width predicts extinction from climate change and vulnerability of tropical species. Global Change Biology, 29(3), 618–630.

Hanski, I. (1994). A practical model of metapopulation dynamics. The Journal of Animal Ecology, 63(1), 151–162.

Hanski, I. (1998). Metapopulation dynamics. Nature, 396(6706), 41–49.

Hanski, I., Moilanen, A., & Gyllenberg, M. (1996). Minimum viable metapopulation size. The American Naturalist, 147(4), 527–541.

Hanski, I., & Ovaskainen, O. (2000). The metapopulation capacity of a fragmented landscape. Nature, 404(6779), 755–758.

Hargreaves, A. L., & Eckert, C. G. (2014). Evolution of dispersal and mating systems along geographic gradients: implications for shifting ranges. Functional Ecology, 28(1), 5–21.

Hargreaves, A. L., Samis, K. E., & Eckert, C. G. (2014). Are species’ range limits simply niche limits writ large? A review of transplant experiments beyond the range. The American Naturalist, 183(2), 157–173.

Hartig, F. (2022). DHARMa: Residual Diagnostics for Hierarchical (Multi-Level / Mixed) Regression Models. http://florianhartig.github.io/DHARMa/

Helenbrook, W. D., & Valdez, J. (2025). Role of rivers as geographical barriers in shaping molecular divergence of neotropical primates. Biotropica, 57(3), e70028.

Holt, R. D., & Keitt, T. H. (2000). Alternative causes for range limits: a metapopulation perspective. Ecology Letters, 3(1), 41–47.

Janitza, S., & Hornung, R. (2018). On the overestimation of random forest’s out-of-bag error. PloS One, 13(8), e0201904.

Jeffries, M. (2008). The spatial and temporal heterogeneity of macrophyte communities in thirty small, temporary ponds over a period of ten years. Ecography, 31(6), 765–775.

Johst, K., Brandl, R., & Eber, S. (2002). Metapopulation persistence in dynamic landscapes: The role of dispersal distance. Oikos, 98(2), 263–270.

Komonen, A. (2007). Are we conserving peripheral populations? An analysis of range structure of longhorn beetles (Coleoptera: Cerambycidae) in Finland. Journal of Insect Conservation, 11(3), 281–285.

Lee-Yaw, J. A., Kharouba, H. M., Bontrager, M., Mahony, C., Csergő, A. M., Noreen, A. M. E., Li, Q., Schuster, R., & Angert, A. L. (2016). A synthesis of transplant experiments and ecological niche models suggests that range limits are often niche limits. Ecology Letters, 19(6), 710–722.

Lennon, J. J., John R. G. Turner, & Connell, D. (1997). A metapopulation model of species boundaries. Oikos, 78(3), 486–502.

Lesica, P., & Allendorf, F. W. (1995). When are peripheral populations valuable for conservation? Conservation Biology: The Journal of the Society for Conservation Biology, 9(4), 753–760.

Littlefield, C. E., Krosby, M., Michalak, J. L., & Lawler, J. J. (2019). Connectivity for species on the move: supporting climate-driven range shifts. Frontiers in Ecology and the Environment, 17(5), 270–278.

Lüdecke, D., Ben-Shachar, M. S., Patil, I., Waggoner, P., & Makowski, D. (2021). performance: An R Package for Assessment, Comparison and Testing of Statistical Models. In Journal of Open Source Software (Vol. 6, Issue 60, p. 3139). 10.21105/joss.03139

Malavasi, M., Bartak, V., Carranza, M. L., Simova, P., & Acosta, A. T. R. (2018). Landscape pattern and plant biodiversity in Mediterranean coastal dune ecosystems: Do habitat loss and fragmentation really matter? Journal of Biogeography, 45(6), 1367–1377.

McLachlan, A. (1991). Ecology of coastal dune fauna. Journal of Arid Environments, 21(2), 229–243.

McPeek, M. A., & Holt, R. D. (1992). The evolution of dispersal in spatially and temporally varying environments. The American Naturalist, 140(6), 1010–1027.

Merriam, C. H. (1894). Laws of temperature control of the geographic distribution of terrestrial animals and plants.

Mestre, F., Pita, R., Mira, A., & Beja, P. (2020). Species traits, patch turnover and successional dynamics: When does intermediate disturbance favour metapopulation occupancy? BMC Ecology, 20(1), 2.

Moilanen, A., & Hanski, I. (1998). Metapopulation dynamics: Effects of habitat quality and landscape structure. Ecology, 79(7), 2503–2515.

Pellet, J., Fleishman, E., Dobkin, D. S., Gander, A., & Murphy, D. D. (2007). An empirical evaluation of the area and isolation paradigm of metapopulation dynamics. Biological Conservation, 136(3), 483–495.

Pinto, S. M., & MacDougall, A. S. (2010). Dispersal limitation and environmental structure interact to restrict the occupation of optimal habitat. The American Naturalist, 175(6), 675–686.

Pironon, S., Papuga, G., Villellas, J., Angert, A. L., García, M. B., & Thompson, J. D. (2017). Geographic variation in genetic and demographic performance: new insights from an old biogeographical paradigm. Biological Reviews of the Cambridge Philosophical Society, 92(4), 1877–1909.

Posit team. (2025). RStudio: Integrated Development Environment for R. Posit Software, PBC. http://www.posit.co/

R Core Team. (2025). R: A Language and Environment for Statistical Computing. R Foundation for Statistical Computing. https://www.R-project.org/

Reed, D. C., Raimondi, P. T., Carr, M. H., & Goldwasser, L. (2000). The role of dispersal and disturbance in determining spatial heterogeneity in sedentary organisms. Ecology, 81(7), 2011–2026.

Rehm, E. M., Olivas, P., Stroud, J., & Feeley, K. J. (2015). Losing your edge: climate change and the conservation value of range-edge populations. Ecology and Evolution, 5(19), 4315–4326.

Reigada, C., Schreiber, S. J., Altermatt, F., & Holyoak, M. (2015). Metapopulation dynamics on ephemeral patches. The American Naturalist, 185(2), 183–195.

Sahlin, E., & Schroeder, L. M. (2010). Importance of habitat patch size for occupancy and density of aspen-associated saproxylic beetles. Biodiversity and Conservation, 19(5), 1325–1339.

Samis, K. E., & Eckert, C. G. (2007). Testing the abundant center model using range-wide demographic surveys of two coastal dune plants. Ecology, 88(7), 1747–1758.

Samis, K. E., & Eckert, C. G. (2009). Ecological correlates of fitness across the northern geographic range limit of a Pacific Coast dune plant. Ecology, 90(11), 3051–3061.

Samis, K. E., López-Villalobos, A., & Eckert, C. G. (2016). Strong genetic differentiation but not local adaptation toward the range limit of a coastal dune plant. Evolution, 70(11), 2520–2536.

Sawchik, J., Dufrêne, M., & Lebrun, P. (2003). Estimation of habitat quality based on plant community, and effects of isolation in a network of butterfly habitat patches. Acta Oecologica, 24(1), 25–33.

Schooley, R. L., & Branch, L. C. (2009). Enhancing the area-isolation paradigm: Habitat heterogeneity and metapopulation dynamics of a rare wetland mammal. Ecological Applications, 19(7), 1708–1722.

Sexton, J. P., McIntyre, P. J., Angert, A. L., & Rice, K. J. (2009). Evolution and ecology of species range limits. Annual Review of Ecology, Evolution, and Systematics, 40(1), 415–436.

Shake, C. S., Moorman, C. E., Riddle, J. D., & Burchell, M. R. (2012). Influence of patch size and shape on occupancy by shrubland birds. The Condor, 114(2), 268–278.

Thomas, J. A., Rose, R. J., Clarke, R. T., Thomas, C. D., & Webb, N. R. (1999). Intraspecific variation in habitat availability among ectothermic animals near their climatic limits and their centres of range. Functional Ecology, 13(s1), 55–64.

Tomiolo, S., & Ward, D. (2018). Species migrations and range shifts: A synthesis of causes and consequences. Perspectives in Plant Ecology, Evolution and Systematics, 33, 62–77.

Travis, J. M. J., & Dytham, C. (1999). Habitat persistence, habitat availability and the evolution of dispersal. Proceedings of the Royal Society of London. Series B: Biological Sciences, 266(1420), 723–728.

Van Natto, A. C., & Eckert, C. G. (2022). Genetic and conservation significance of populations at the polar vs. equatorial range limits of the Pacific coastal dune endemic Abronia umbellata (Nyctaginaceae). Conservation Genetics, 23(2), 255–269.

Venable, D. L. (2007). Bet hedging in a guild of desert annuals. Ecology, 88(5), 1086–1090.

Vuilleumier, S., Wilcox, C., Cairns, B. J., & Possingham, H. P. (2007). How patch configuration affects the impact of disturbances on metapopulation persistence. Theoretical Population Biology, 72(1), 77–85.

Walton, J. (2008). The land of Big Sur: Conservation on the California coast. California History, 85(1), 44–64.

White, E. R., & Smith, A. T. (2018). The role of spatial structure in the collapse of regional metapopulations. Ecology, 99(12), 2815–2822.

Wilcox, C., Cairns, B. J., & Possingham, H. P. (2006). The role of habitat disturbance and recovery in metapopulation persistence. Ecology, 87(4), 855–863.

