## Supporting Information for "Spatial and temporal habitat availability declines towards and beyond the geographic range limit of a coastal dune endemic"

MANUSCRIPT TITLE

APPENDIX S1

**Supplementary Figures & Tables**

**Table S1.** Binomial GLM analyses of the effects of patch characteristics and range position on patch occupancy by coastal dune endemic *Camissoniopsis cheiranthifolia* in 2019 and 2022. All predictors are scaled for comparability, and all coefficients are in log-odds. Significant predictors are bolded.

| Response | *n* | Predictor | df | *b* | χ*^2^* | *P* |
| --- | --- | --- | --- | --- | --- | --- |
| 2019 occupancy | 1293 | **Patch size** | **1** | **0.92** | **155.37** | **<0.0001** |
|  |  | **Patch isolation** | **1** | **–0.23** | **11.30** | **0.00078** |
|  |  | **Patch stability** | **1** | **0.74** | **14.03** | **0.00018** |
|  |  | **Distance to range limit** | **1** | **0.41** | **38.91** | **<0.0001** |
| 2022 occupancy | 2088 | **Patch size** | **1** | **0.80** | **211.88** | **<0.0001** |
|  |  | Patch isolation | 1 | –0.11 | 3.11 | 0.078 |
|  |  | **Patch stability** | **1** | **1.28** | **100.89** | **<0.0001** |
|  |  | **Distance to range limit** | **1** | **0.54** | **89.53** | **<0.0001** |

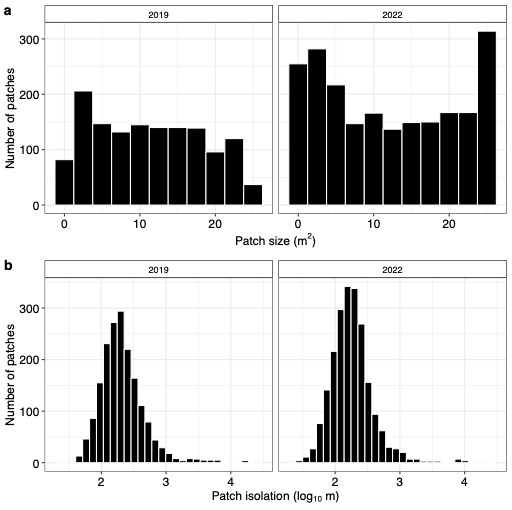

*Figure S1.* The distribution of patch sizes (m^2^; panel a), excluding patches that were entirely unsuitable, and patch isolation (log_10_ m; panel b) in both 2019 (left) and 2022 (right).
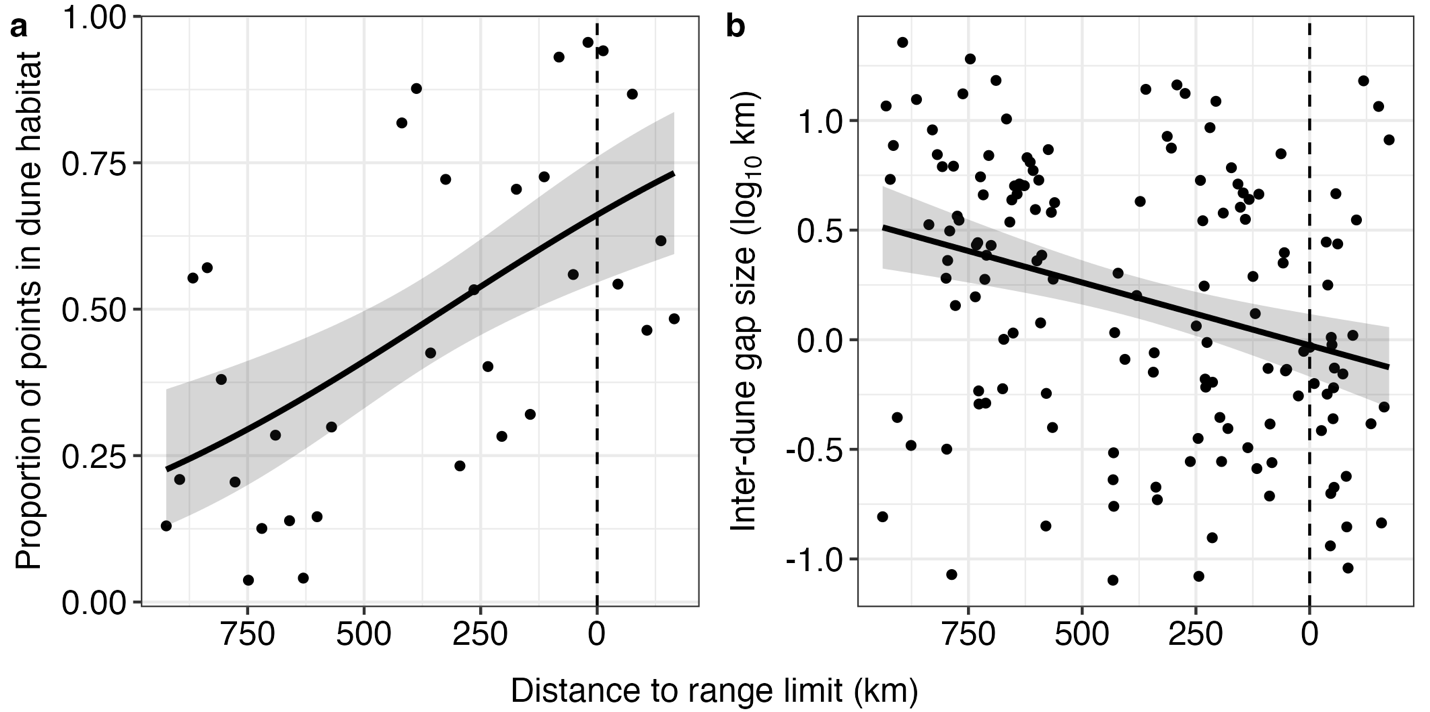
*Figure S2.* (a) The proportion of points occurring within coastal dune habitat increase towards the range limit of *Camissoniopsis cheiranthifolia* after excluding all bins of 2500 points with zero dune habitat and (b) gap sizes between adjacent coastal dune sites decline towards the range limit after excluding the large gap at the Lost Coast. Grey ribbons are 95% confidence intervals. The dashed lines indicate the northern range limit.

APPENDIX S2

Random forest model performance

Table S2. List of species observed throughout the 2022 survey of 4023 randomly distributed 5m x 5m plots were recorded, ordered by the % increase in mean standard error (MSE) in the random forest model used to predict *Camissoniopsis cheiranthifolia* occupancy. Columns indicate the number of plots where the species was observed (Count) and the survey-wide frequency (Freq.), as well as the maximum and minimum latitude where the species was observed.

| Rank order | % increase in MSE | Species | Count | Freq. | Max. lat. (˚N) | Min. lat. (˚S) |
| --- | --- | --- | --- | --- | --- | --- |
| 1 | 46.63 | *Artemisia pycnocephala* | 164 | 4.08 | 41.87 | 37.80 |
| 2 | 40.77 | *Ammophila arenaria* | 2253 | 56.00 | 45.22 | 37.81 |
| 3 | 39.94 | *Calystegia soldanella* | 383 | 9.52 | 45.17 | 37.81 |
| 4 | 38.17 | *Solidago spathulata* | 161 | 4.00 | 45.18 | 39.49 |
| 5 | 32.60 | *Abronia latifolia* | 129 | 3.21 | 44.43 | 37.81 |
| 6 | 30.78 | *Polygonum paronychia* | 260 | 6.46 | 45.22 | 38.11 |
| 7 | 27.64 | *Lupinus littoralis* | 202 | 5.02 | 45.22 | 38.02 |
| 8 | 25.79 | *Fragaria chiloensis* | 222 | 5.52 | 45.22 | 37.80 |
| 9 | 24.31 | *Lathyrus japonicus* | 358 | 8.90 | 45.22 | 40.99 |
| 10 | 23.99 | *Baccharis pilularis* | 454 | 11.29 | 44.38 | 37.81 |
| 11 | 23.77 | *Abronia umbellata* | 13 | 0.32 | 43.37 | 37.89 |
| 12 | 22.91 | *Cardionema ramosissimum* | 81 | 2.01 | 45.13 | 38.10 |
| 13 | 19.78 | *Ambrosia chamissonis* | 132 | 3.28 | 42.79 | 37.80 |
| 14 | 18.70 | *Anaphalis margaritaceae* | 155 | 3.85 | 45.18 | 38.03 |
| 15 | 17.64 | *Rumex acetosella* | 308 | 7.66 | 45.22 | 38.10 |
| 16 | 17.44 | *Lathyrus littoralis* | 142 | 3.53 | 45.18 | 38.11 |
| 17 | 17.40 | *Rubus ursinus* | 63 | 1.57 | 45.01 | 38.02 |
| 18 | 16.17 | *Gaultheria shallon* | 87 | 2.16 | 45.01 | 41.20 |
| 19 | 15.16 | *Carpobrotus chilensis* | 38 | 0.94 | 42.74 | 37.90 |
| 20 | 14.13 | *Tanacetum bipinnatum* | 50 | 1.24 | 45.00 | 38.05 |
| 21 | 13.91 | *Cytisus scoparius* | 139 | 3.46 | 45.22 | 39.53 |
| 22 | 11.54 | *Picea sitchensis* | 116 | 2.88 | 45.19 | 40.75 |
| 23 | 11.47 | *Heterotheca sessiliflora* | 10 | 0.25 | 39.55 | 39.48 |
| 24 | 11.39 | *Achillea millefolium* | 206 | 5.12 | 45.19 | 38.10 |
| 25 | 11.07 | *Ericameria ericoides* | 12 | 0.30 | 38.31 | 38.10 |
| 26 | 11.03 | *Pseudognaphalium luteoalbum* | 82 | 2.04 | 43.09 | 38.03 |
| 27 | 9.23 | *Carex obnupta* | 39 | 0.97 | 44.46 | 38.08 |
| 28 | 8.01 | *Daucus pusillus* | 44 | 1.09 | 43.47 | 40.90 |
| 29 | 7.84 | *Senecio glomeratus* | 18 | 0.45 | 43.63 | 38.97 |
| 30 | 7.72 | *Acmispon heermannii* | 6 | 0.15 | 38.32 | 38.11 |
| 31 | 7.52 | *Morella californica* | 36 | 0.89 | 44.99 | 41.35 |
| 32 | 7.31 | *Gamochaeta ustulata* | 97 | 2.41 | 45.22 | 38.32 |
| 33 | 6.46 | *Pinus contorta* | 113 | 2.81 | 45.19 | 38.03 |
| 34 | 5.90 | *Ulex europaeus* | 17 | 0.42 | 44.96 | 42.79 |
| 35 | 5.86 | *Erigeron glaucus* | 41 | 1.02 | 43.35 | 37.89 |
| 36 | 5.09 | *Lupinus arboreus* | 146 | 3.63 | 42.28 | 38.03 |
| 37 | 5.08 | *Elymus mollis* | 331 | 8.23 | 45.22 | 37.90 |
| 38 | 4.83 | *Raphanus sativus* | 4 | 0.10 | 44.93 | 39.43 |
| 39 | 4.71 | *Hirschfeldia incana* | 2 | 0.05 | 37.89 | 37.89 |
| 40 | 4.22 | *Pentagramma triangularis* | 42 | 1.04 | 45.11 | 40.83 |
| 41 | 4.19 | *Clinopodium douglasii* | 4 | 0.10 | 42.26 | 41.89 |
| 42 | 3.73 | *Glehnia littoralis* | 25 | 0.62 | 42.97 | 41.14 |
| 43 | 3.55 | *Bromus diandrus* | 6 | 0.15 | 43.96 | 42.09 |
| 44 | 3.52 | *Vaccinium ovatum* | 30 | 0.75 | 44.92 | 42.45 |
| 45 | 3.02 | *Holcus lanatus* | 156 | 3.88 | 45.12 | 38.02 |
| 46 | 2.92 | *Grindelia stricta* | 16 | 0.40 | 41.81 | 38.02 |
| 47 | 2.88 | *Plantago coronopus* | 9 | 0.22 | 43.14 | 41.28 |
| 48 | 2.66 | *Bellardia viscosa* | 4 | 0.10 | 44.46 | 40.98 |
| 49 | 2.29 | *Sonchus asper* | 50 | 1.24 | 45.19 | 38.96 |
| 50 | 2.25 | *Digitalis purpurea* | 4 | 0.10 | 45.12 | 42.26 |
| 51 | 2.24 | *Raphanus raphanistrum* | 7 | 0.17 | 42.74 | 39.43 |
| 52 | 1.95 | *Plantago lanceolata* | 49 | 1.22 | 44.64 | 38.11 |
| 53 | 1.95 | *Cupressus macrocarpa* | 5 | 0.12 | 40.67 | 38.33 |
| 54 | 1.83 | *Cortaderia selloana* | 5 | 0.12 | 41.02 | 40.84 |
| 55 | 1.63 | *Dactylis glomerata* | 17 | 0.42 | 44.40 | 42.35 |
| 56 | 1.57 | *Diplacus aurantiacus* | 2 | 0.05 | 38.45 | 38.03 |
| 57 | 1.46 | *Jacobaea vulgaris* | 9 | 0.22 | 45.19 | 42.82 |
| 58 | 1.09 | *Erigeron canadensis* | 8 | 0.20 | 42.45 | 39.50 |
| 59 | 1.04 | *Genista monspessulana* | 5 | 0.12 | 42.10 | 41.00 |
| 60 | 1.02 | *Acmispon americanus* | 3 | 0.07 | 41.20 | 41.20 |
| 61 | 0.83 | *Salicornia pacifica* | 3 | 0.07 | 40.71 | 40.58 |
| 62 | 0.46 | *Polycarpon tetraphyllum* | 34 | 0.85 | 42.45 | 40.61 |
| 63 | 0.35 | *Claytonia perfoliata* | 3 | 0.07 | 42.42 | 42.34 |
| 64 | 0.00 | *Artemisia douglasiana* | 2 | 0.05 | 42.82 | 42.04 |
| 64 | 0.00 | *Atriplex leucophylla* | 1 | 0.02 | 38.99 | 38.99 |
| 64 | 0.00 | *Atriplex prostrata* | 1 | 0.02 | 40.62 | 40.62 |
| 64 | 0.00 | *Bromus carinatus* | 2 | 0.05 | 43.49 | 42.26 |
| 64 | 0.00 | *Cakile edentula* | 4 | 0.10 | 45.12 | 42.39 |
| 64 | 0.00 | *Cirsium vulgare* | 1 | 0.02 | 43.54 | 43.54 |
| 64 | 0.00 | *Comarum palustre* | 1 | 0.02 | 43.48 | 43.48 |
| 64 | 0.00 | *Cotoneaster sp.* | 1 | 0.02 | 42.05 | 42.05 |
| 64 | 0.00 | *Cyperus eragrostis* | 1 | 0.02 | 41.43 | 41.43 |
| 64 | 0.00 | *Hordeum murinum* | 1 | 0.02 | 42.39 | 42.39 |
| 64 | 0.00 | *Koeleria macrantha* | 1 | 0.02 | 38.97 | 38.97 |
| 64 | 0.00 | *Logfia gallica* | 1 | 0.02 | 42.27 | 42.27 |
| 64 | 0.00 | *Lolium perenne* | 1 | 0.02 | 42.43 | 42.43 |
| 64 | 0.00 | *Lupinus bicolor* | 1 | 0.02 | 42.79 | 42.79 |
| 64 | 0.00 | *Lupinus chamissonis* | 1 | 0.02 | 38.11 | 38.11 |
| 64 | 0.00 | *Lysichiton americanus* | 1 | 0.02 | 43.14 | 43.14 |
| 64 | 0.00 | *Phalaris arundinacea* | 1 | 0.02 | 43.09 | 43.09 |
| 64 | 0.00 | *Polystichum imbricans* | 1 | 0.02 | 39.66 | 39.66 |
| 64 | 0.00 | *Poa macrantha* | 1 | 0.02 | 41.18 | 41.18 |
| 64 | 0.00 | *Polystichum munitum* | 4 | 0.10 | 43.96 | 43.51 |
| 64 | 0.00 | *Scrophularia californica* | 1 | 0.02 | 40.91 | 40.91 |
| 64 | 0.00 | *Schoenoplectus pungens* | 1 | 0.02 | 42.10 | 42.10 |
| 64 | 0.00 | *Sequoia sempervirens* | 1 | 0.02 | 41.93 | 41.93 |
| 64 | 0.00 | *Spergularia macrotheca* | 1 | 0.02 | 40.63 | 40.63 |
| 64 | 0.00 | *Visia hirsuta* | 1 | 0.02 | 42.26 | 42.26 |
| 64 | 0.00 | *Vicia villosa* | 1 | 0.02 | 42.04 | 42.04 |
| 90 | -0.12 | *Lotus corniculatus* | 23 | 0.57 | 41.39 | 38.02 |
| 91 | -0.33 | *Potentilla anserina* | 9 | 0.22 | 45.01 | 41.72 |
| 92 | -0.37 | *Mentha pulegium* | 4 | 0.10 | 41.43 | 41.41 |
| 93 | -0.39 | *Visia sativa* | 29 | 0.72 | 43.66 | 39.02 |
| 94 | -0.43 | *Equisetum telmateia* | 3 | 0.07 | 43.19 | 39.66 |
| 95 | -0.56 | *Typha latifolia* | 2 | 0.05 | 41.18 | 39.68 |
| 96 | -0.57 | *Lupinus tidestromii* | 2 | 0.05 | 38.11 | 38.11 |
| 97 | -0.89 | *Lonicera involucrata* | 11 | 0.27 | 44.47 | 41.00 |
| 98 | -1.00 | *Vicia tetrasperma* | 3 | 0.07 | 41.29 | 40.94 |
| 99 | -1.07 | *Heracleum maximum* | 4 | 0.10 | 42.82 | 41.93 |
| 100 | -1.09 | *Briza maxima* | 87 | 2.16 | 44.01 | 39.49 |
| 101 | -1.22 | *Plantago erecta* | 9 | 0.22 | 43.14 | 40.89 |
| 102 | -1.42 | *Pteridium aquilinum* | 8 | 0.20 | 44.41 | 38.02 |
| 103 | -1.52 | *Anthoxanthum odoratum* | 33 | 0.82 | 45.11 | 39.30 |
| 104 | -1.67 | *Geranium dissectum* | 3 | 0.07 | 42.26 | 40.94 |
| 105 | -1.74 | *Senecio minimus* | 16 | 0.40 | 45.18 | 42.82 |
| 106 | -1.84 | *Silene gallica* | 26 | 0.65 | 42.85 | 41.23 |
| 107 | -2.01 | *Polypodium glycyrrhiza* | 2 | 0.05 | 43.51 | 43.50 |
| 108 | -2.03 | *Dudleya farinosa* | 3 | 0.07 | 38.04 | 38.04 |
| 109 | -2.04 | *Plantago maritima* | 5 | 0.12 | 43.11 | 39.19 |
| 110 | -2.27 | *Alnus rubra* | 5 | 0.12 | 44.52 | 41.01 |
| 111 | -2.28 | *Bromus hordeaceus* | 6 | 0.15 | 43.45 | 39.00 |
| 112 | -2.35 | *Eriophyllum staechadifolium* | 5 | 0.12 | 39.66 | 38.11 |
| 113 | -2.41 | *Ammophila breviligulata* | 4 | 0.10 | 43.71 | 42.10 |
| 114 | -2.47 | *Calamagrostis epigejos* | 3 | 0.07 | 41.27 | 41.01 |
| 115 | -2.73 | *Medicago polymorpha* | 3 | 0.07 | 42.43 | 41.28 |
| 116 | -3.35 | *Cerastium glomeratum* | 16 | 0.40 | 45.12 | 40.90 |
| 117 | -3.41 | *Cynosurus echinatus* | 42 | 1.04 | 45.19 | 38.76 |
| 118 | -3.69 | *Pseudognaphalium stramineum* | 2 | 0.05 | 42.00 | 41.21 |
| 119 | -5.31 | *Galium aparine* | 49 | 1.22 | 45.19 | 40.90 |
| 120 | -6.03 | *Cakile maritima* | 120 | 2.98 | 45.00 | 37.89 |

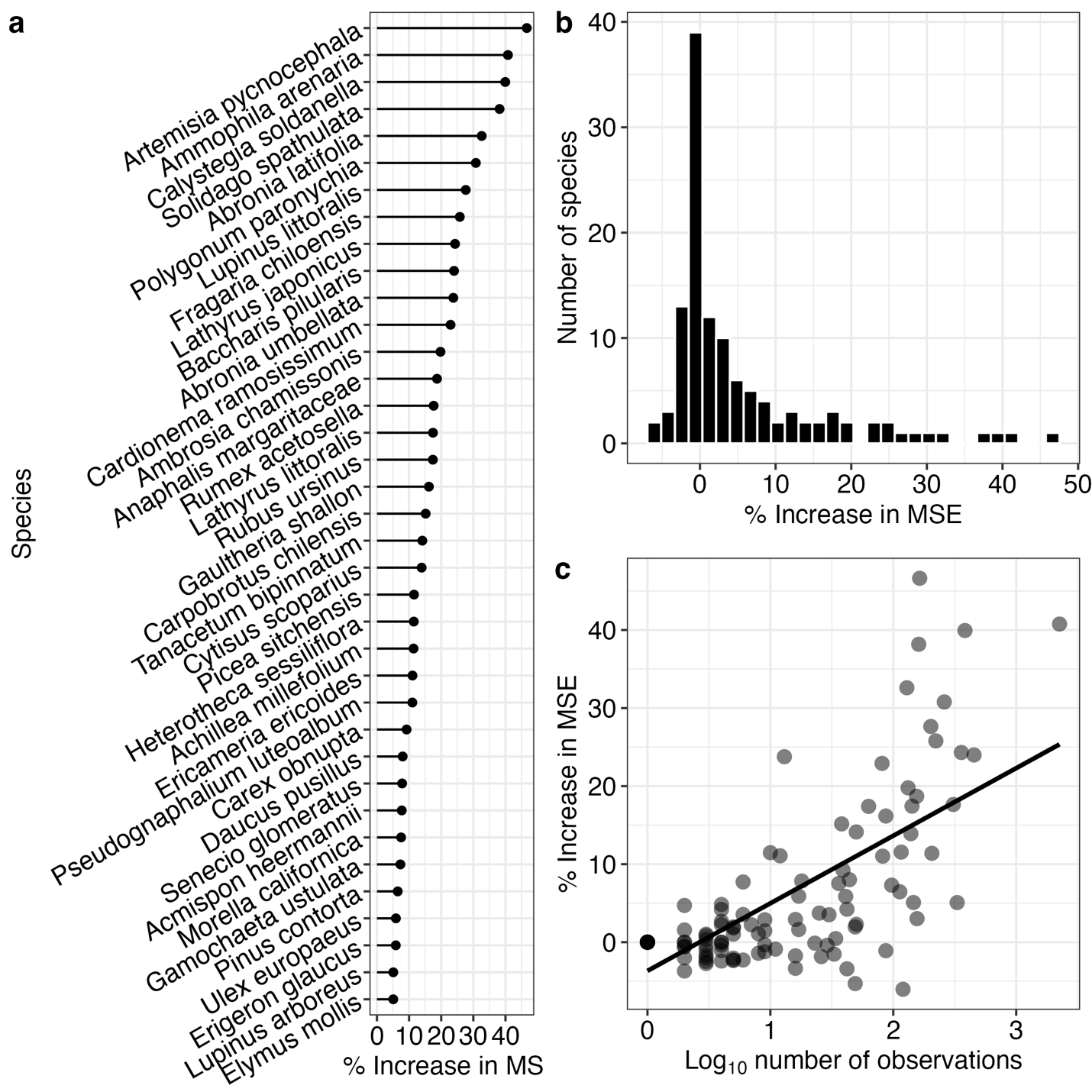

*Figure* *S3.* (a) Variable importance plots of species used in the random forest model predicting the occupancy of *Camissoniopsis cheiranthifolia*, where all species with a % increase in mean standard error (MSE) > 5% are ranked by MSE. (b) The distribution of the percent increase in MSE for all species observed in the survey. (c) The percent increase in MSE of species in the RF model is positively correlated with the log_10_-transformed number of times the species was observed (Gaussian generalized linear model, *n* = 120, *b* = 8.66, χ*^2^* = 107.58, df = 1, *P* < 0.0001).
